# Mapping the Developmental Trajectory of Functional Brain Networks in Early Infancy: Insights into Typical Maturation^*^

**DOI:** 10.1101/2025.11.25.690500

**Authors:** Masoud Seraji, Sarah Shultz, Qiang Li, Zening Fu, Armin Iraji, Vince D. Calhoun

**Affiliations:** Tri-institutional Center for Translational Research in Neuroimaging and Data Science (TReNDS): Georgia State University, Georgia Institute of Technology, Emory University, Atlanta, GA 30303, USA; The school of psychology, University of Texas at Austin, Austin, TX 78712, USA; Department of Pediatrics and Marcus Autism Center, Emory Univeristy, Atlanta, GA 30329USA

## Abstract

Early infancy is a crucial period for brain development, during which fundamental functional and structural frameworks are established. Understanding the maturation of large-scale brain networks during this stage is essential for characterizing normative neurodevelopment and identifying potential deviations linked to neurodevelopmental disorders. In this study, we investigated developmental changes in the spatial organization of functional brain networks in infants using a longitudinal resting-state fMRI dataset comprising 137 scans from 74 low-likelihood developing infants aged 0–6 months. We applied independent component analysis to extract large-scale brain networks and utilized advanced spatial metrics, including network-averaged spatial similarity (NASS) to assess alignment with group-level patterns, network strength to quantify neural engagement based on voxel intensities, and network size to examine spatial distribution. Our findings reveal significant age-related increases in NASS across multiple networks, indicating greater consistency in functional organization over time. Additionally, most networks demonstrated increased network strength, reflecting heightened neural involvement, while network size exhibited distinct developmental trajectories, with some networks expanding and others remaining stable. These results highlight the dynamic evolution of functional brain architecture during early infancy, providing critical insights into neurodevelopmental processes.

**Clinical Relevance:** This study provides critical insights into early brain network development, which is essential for identifying biomarkers of neurodevelopmental disorders such as autism and schizophrenia. By mapping typical maturation patterns using advanced spatial metrics, our findings offer a foundation for early detection of atypical development. Deviations in network organization and strength could serve as early indicators, supporting neuroimaging-based screening and intervention strategies to optimize neurodevelopmental outcomes.

## I. Introduction

The first few months after birth is considered a crucial timeframe in which the fundamental aspects of the brain’s functional and structural framework are established [1], [2]. Additionally, there is a growing understanding that developmental disorders and psychopathology often originate in infancy [1]. This period is recognized as a crucial factor in the predisposition to conditions such as autism and schizophrenia [3]. These disorders typically emerge with symptoms during childhood and later developmental stages, influenced by the intricate dynamics of early brain development [1], [4], [5]. This highlights the significance of understanding the early stages of brain development to potentially mitigate or manage the onset and progression of these disorders. Moreover, functional brain networks have been linked to various behaviors and developmental trajectories [5]. However, despite extensive research on infant development, the specific patterns of change across large-scale brain networks in the first few months of life have not been fully understood [5]. Therefore, there is a significant need to track and study the developmental trajectory of the brain networks during this pivotal period. Previous works has used functional connectivity (FC) [6], or its network analog, functional network connectivity (FNC), to study developmental changes in late childhood and adolescence [4], [7], [8]. However, there is a notable gap in our understanding when it comes to describing spatial maps developments during infancy. While there have been observations of precursors to mature functional networks during gestation and in both premature [9], [10] and full-term infants at birth [11], [12], most studies have focused on examining developmental differences in FC over relatively lengthy periods (e.g., from infancy to adulthood). In the context of extracting large-scale brain networks, independent component analysis (ICA) [13] stands out as a widely utilized and highly effective technique. Its utility lies in its ability to identify common brain networks, thus deepening our understanding of brain function [14], [15], [16]. While our methodology closely aligns with many studies employing ICA on fMRI data in adults, which typically involves group-level ICA followed by back-reconstruction, we have enhanced our back-reconstruction process compared to previous infant studies. This improvement involves the adoption of group information-guided ICA (GIG-ICA) [17], which enhances spatial consistency across subjects.

In our study, we investigated developmental changes in the spatial organization of functional brain networks in infants using a comprehensive set of spatial metrics. We employed network-averaged spatial similarity (NASS) to assess how individual spatial maps align with the group-level network, network strength to measure neural activity, and network size to examine spatial distribution. By integrating these metrics, we effectively captured age-related variations in large-scale brain networks, providing insights into the refinement and maturation of brain networks during early development. Our findings contribute to understanding normative neurodevelopment and may serve as a foundation for identifying deviations associated with atypical brain maturation.

## II. Methods

### A. Imaging Dataset

We collected a longitudinal dataset of 137 resting-state fMRI scans from 74 low-likelihood developing infants (43 males, 31 females) aged 0–3 months, with up to three scans per infant. None of the infants had a family history of autism spectrum disorder or clinical concerns [18], [19]. Corrected age, calculated based on a standard gestational age of 40 weeks, was used for developmental analyses to ensure accuracy. All parents provided informed consent, and the study was approved by Emory University.

### B. Preprocessing

We preprocessed the data by discarding the initial 16 volumes to stabilize magnetization and corrected head motion using FSL’s mcflirt function. Distortion in multi-band rs-fMRI data was corrected using single-band data with AP and PA phase encoding to estimate susceptibility-induced off-resonance fields, followed by slice timing correction. A two-step normalization process brought the data into a common MNI space: first aligning to a 3-month T1 template from the UNC/UMN Baby Connectome Project using affine transformation, and then to MNI space using an EPI template. The 3-month template was selected for consistency across participants, and additional control analyses using age-specific templates confirmed the robustness of this choice. Finally, we smoothed the normalized images with a 6 mm Gaussian kernel.

### C. Data Analysis

We conducted group ICA using the GIFT toolbox to analyze fMRI scans, starting with subject-specific PCA to standardize data, reduce noise, and retain 30 principal components (PCs) per subject, capturing maximum variance. These PCs were combined across subjects and further analyzed with group-level PCA to identify commonalities. We selected the top 20 group-level PCs and performed group ICA with a model order of 20, optimized to capture large-scale brain networks. The Infomax algorithm was run 100 times with random initialization and bootstrapping, ensuring reliability through ICASSO quality index (IQ), with components selected based on stability (IQ > 0.80), overlap with gray matter, and low similarity to artifacts. Finally, the GIG-ICA technique estimated subject-specific large-scale brain networks.

### D. Spatial Metrics

of the network. We developed several metrics, including network-averaged spatial similarity, network strength, and network size, to investigate developmental changes in the spatial organization of brain networks, as detailed below.

#### Network Strength

Network strength represents the average intensity of all voxels within a network, serving as a measure of the network’s functional engagement. Higher network strength values indicate greater voxel involvement. To calculate this metric, we transformed voxel-wise intensities to Z-scores by subtracting the mean and dividing by the standard deviation across all voxels. A threshold of Z > 1.96 (p = 0.05) was applied to include only significantly contributing voxels, and network strength was then computed as the average intensity of these selected voxels per scan.

#### Network Size

Network size quantifies the spatial extent of a network, providing insight into the expansion or contraction of brain networks over time. Variations in network size reflect changes in the number of voxels contributing to a specific network, indicating fluctuations in its spatial organization. To compute network size, we used the same masking approach to identify voxels with Z-scores above 1.96 (p = 0.05) within each network. We then counted the number of voxels meeting this criterion to derive a value representing the size of the network. This metric captures dynamic changes in network architecture over developmental stages.

#### Network-averaged Spatial Similarity (NASS)

We introduced the metric “network-averaged spatial similarity” (NASS) to evaluate the degree of alignment between individual participant-specific spatial maps and the group-level network map. The group-level map captures the shared spatial organization of brain networks across all participants, representing common patterns. To compute NASS, we correlated each participant’s individual spatial map with the corresponding group-level network, yielding a NASS value for each network in every scan. Higher NASS values indicate a strong alignment between an individual’s network and the group-level spatial pattern, reflecting greater network homogeneity. In contrast, lower NASS values suggest increased deviation from the group-level structure, highlighting more pronounced individual-specific variations in the spatial organization

### E. Statistical Analysis

For the statistical analysis, we utilized a generalized additive model (GAM) [20] to examine the relationships between each metric and the variables of interest. This model incorporated corrected age, sex, scanner type, and head motion as fixed effects, while accounting for individual variability by treating participants as random effects. By accommodating both linear and non-linear associations, this approach enhances the robustness and flexibility of our inferences while effectively controlling for potential confounders. To address the issue of multiple comparisons, we applied false discovery rate (FDR) correction to the p-values, ensuring that significant results reflect true effects rather than random chance.

## III. Results And Discussion

Our results are organized into two primary sections. First, we presented the functional brain networks identified in infants, highlighting their structure and characteristics. Second, we examined the developmental progression of these networks with age, analyzing how our measurements evolved over time to provide insights into their dynamic changes.

### A. Insights into Identified Brain Networks

We conducted ICA on rsfMRI data from 74 subjects, extracting 20 components and successfully identifying 13 distinct brain networks in infants, as illustrated in Fig. 1. These networks include the primary and secondary visual networks, the subcortical network, the cerebellar network, the primary and secondary motor networks, the attention network, the default mode network, the temporal network, the auditory network, and three frontal networks: medial prefrontal cortex (mPFC), dorsolateral prefrontal cortex (dlPFC), and ventrolateral prefrontal cortex (vlPFC). This comprehensive analysis highlights the diversity and organization of functional brain networks during early development.

**Figure 1.**
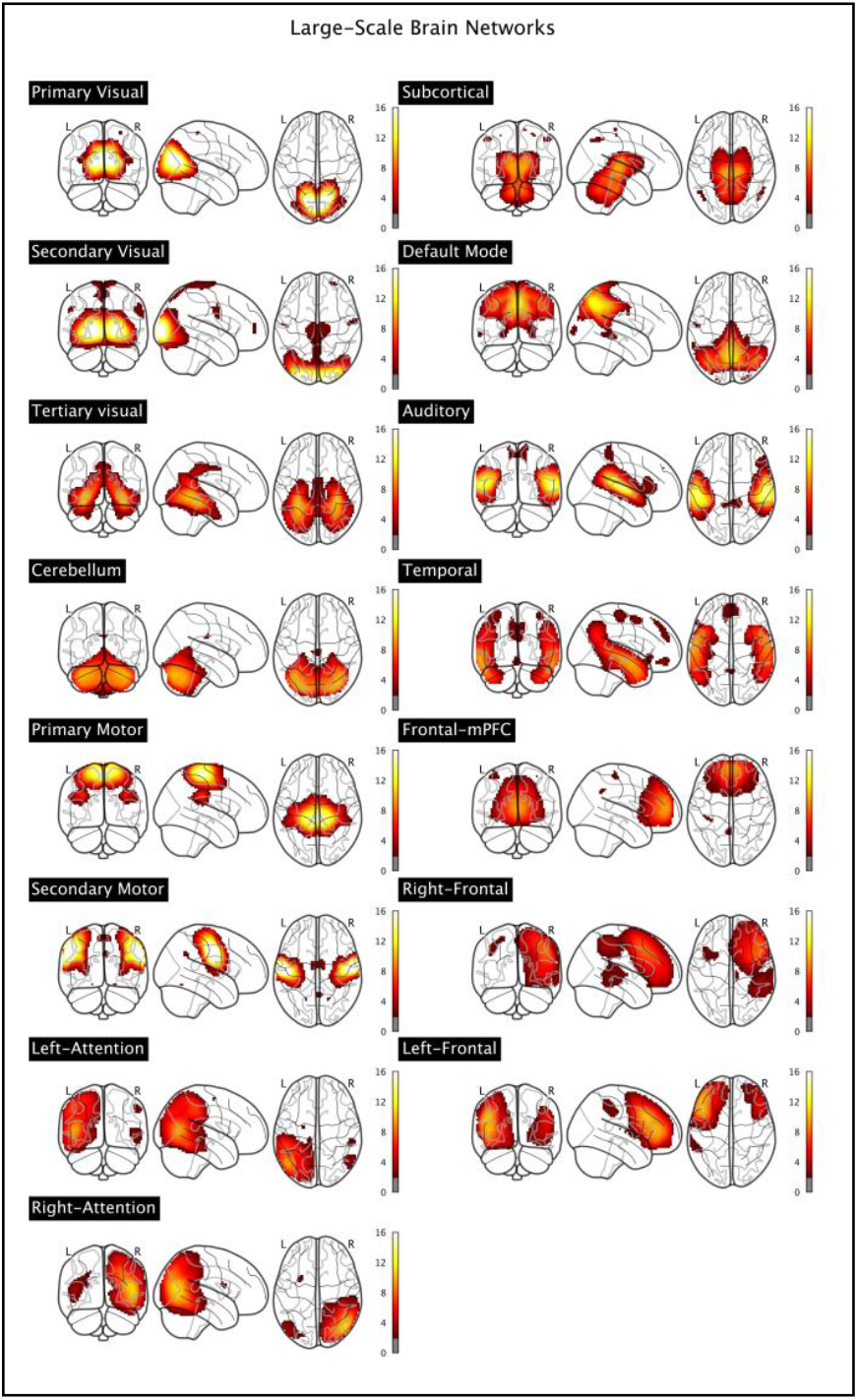
Displaying sagittal, coronal, and axial views of z-scored voxel intensities within the spatial maps of 13 functional brain networks in infants.

### B. Spatial Developmental Patterns in Brain Networks

In this section, we examine the spatial developmental metrics of specific brain networks across different ages. As representative examples, we conducted a detailed assessment of the primary visual, secondary visual, cerebellar, primary motor, secondary motor, attention, and subcortical networks. Figure 2 presents a comprehensive analysis of the temporal evolution of these metrics. Our findings indicate that the Network-Averaged Spatial Similarity of all seven networks except the subcortical network to their respective group-level estimates increases over time (F > 5.45, p < 0.02; Fig. 2), suggesting progressive network refinement with age.

**Figure 2.**
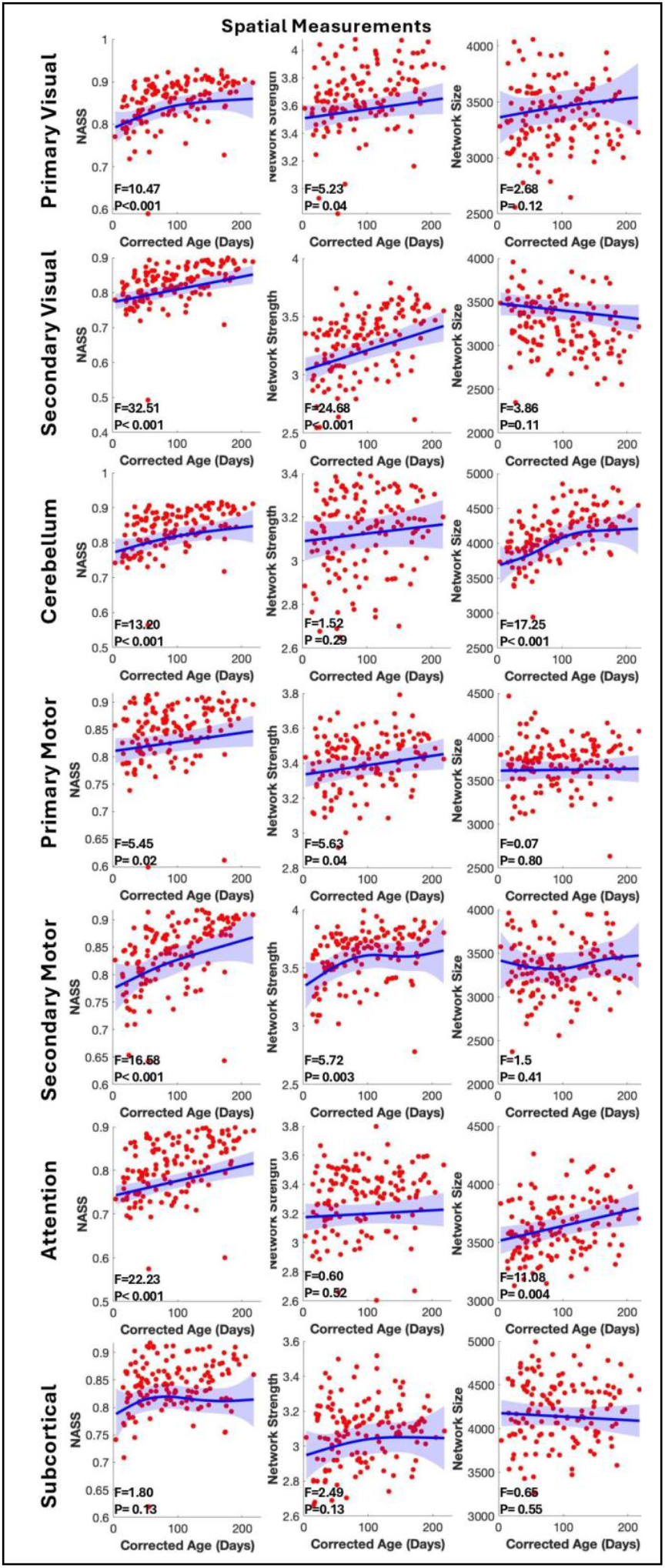
This figure illustrates the variations in spatial metrics across the primary visual, secondary visual, cerebellum, primary motor, secondary motor, attention, and subcortical networks, focusing on network-averaged spatial similarity (NASS), network strength, and network size. Each red dot represents an individual scan obtained from infants across different age groups, while the blue line depicts the overall trend. All p-values are corrected for multiple comparisons using the False Discovery Rate (FDR) method, ensuring statistical robustness.

Additionally, primary visual, secondary visual, primary motor and secondary motor networks showed a significant increase in network strength (F > 5.23, p < 0.04; Fig. 2), indicating heightened activation as infants mature. In terms of network size, the primary visual, secondary visual, primary motor, secondary motor, and subcortical networks exhibited no significant change (F < 3.86, p > 0.11; Fig. 2). Furthermore, age-related analyses revealed substantial growth in the cerebellar and attention networks (F > 11.08, p < 0.004; Fig. 2), emphasizing their increasing role in cognitive and motor development throughout infancy.

Our findings provide a detailed perspective on the spatial changes occurring across various brain networks during early development. Notably, the increasing NASS over time suggests that individual network patterns become more aligned with their averaged group representation as infants mature. This trend may reflect the progressive refinement and stabilization of functional connectivity, reinforcing the robustness of network architecture and hinting at a degree of universality in the spatial organization of brain networks [21]. The consistency observed in NASS may contribute to our broader understanding of normative brain development and functional stability.

In addition to NASS, we observed significant changes in network strength during the first six months of life across various brain networks. This dynamic increase may indicate a critical phase of neurodevelopment, characterized by processes such as myelination, synaptic pruning, and the establishment of neural connections [22]. Alternatively, heightened network strength could reflect greater neural activity and efficiency, suggesting that these networks undergo refinement and optimization during early infancy [23]. These findings align with prior research highlighting the importance of early postnatal development in shaping brain function.

Another key metric we analyzed was network size, which exhibited distinct variation across different brain networks. Specifically, changes were observed in the cerebellum, and attention networks, whereas other networks remained relatively stable [24]. For example, the cerebellum’s network expansion is closely associated with key developmental advancements in motor control, coordination, and cognitive functions. This growth plays a fundamental role in refining fine motor skills, such as reaching and grasping, as well as supporting postural stability and balance, both of which are crucial as infants reach major motor milestones [25]. Additionally, the cerebellum is instrumental in cognitive processes like attention regulation and early learning, reinforcing its dual function in both sensorimotor and higher-order cognitive development [26]. This progressive increase in cerebellar network size highlights its essential role in shaping infants’ ability to engage with their surroundings, facilitating both motor adaptability and cognitive learning. This divergence in spatial dynamics suggests that different neural circuits follow unique developmental trajectories during early life. Networks experiencing significant expansion may be undergoing active synaptogenesis, a crucial process that supports cognitive and sensorimotor functions [27].

### C. Insight Into Visual networks

Fig. 3 illustrates the distribution of NASS and network strength in infants at 1, 3, and 6 months of age. The data reveal a consistent linear increase in these metrics over the six-month period, with one exception: the primary visual network exhibits a nonlinear growth trajectory between birth and three months. Specifically, this network undergoes a rapid increase in NASS within the first three months, followed by a more linear growth pattern from 3 to 6 months. This accelerated early development suggests that the primary visual network is well-established by three months of age.

**Figure 3.**
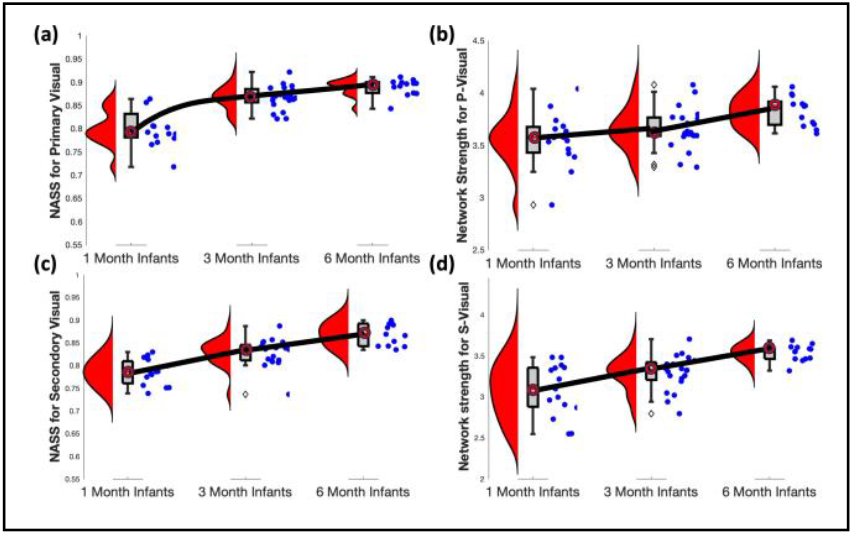
Developmental changes in primary and secondary visual networks over the first six months of infancy. (a) Network-average spatial similarity (NASS) for the primary visual network shows a rapid increase between 1 and 3 months, followed by a stabilization from 3 to 6 months. (b) Network strength of the primary visual network exhibits a more gradual increase over time. (c) NASS for the secondary visual network follows a linear gradual increase. (d) Network strength of the secondary visual network demonstrates a steady increase across all three time points. Boxplots represent data distributions, while violin plots (red) illustrate density estimates. Blue dots indicate individual data points, and black lines represent fitted trends.

The early maturation of visual networks, coupled with the stability in their network size, aligns with previous research showing that these networks are functional at birth. While they continue to develop during the first six months of life, their growth occurs at a slower rate compared to other networks. Nonetheless, visual networks are among the first to achieve an adult-like state, reflecting their early importance in sensory processing and perceptual development [28], [29]. Notably, sensorimotor networks exhibit a more rapid expansion compared to visual networks during early childhood, consistent with previous findings [30]. This trend is reflected in our results, where the primary and secondary motor networks not only demonstrate greater overall strength but also a non-linear rate of increase relative to visual networks.

This differential growth pattern underscores the dynamic nature of neurodevelopment, where visual networks establish early functionality, while sensorimotor networks continue evolving to support complex motor and cognitive functions. Additional research is needed to investigate the underlying mechanisms driving these distinct developmental trajectories and to explore their implications for long-term cognitive and behavioral outcomes.

## IV. Conclusion

This study examined the developmental changes in functional brain networks during early infancy using independent component analysis and advanced spatial metrics. Our findings reveal significant age-related increases in network-averaged spatial similarity, indicating greater alignment with group-level network structures over time. Additionally, most networks exhibited increased network strength, reflecting heightened neural engagement, while network size showed distinct developmental trajectories. These results highlight the dynamic maturation of intrinsic connectivity networks in the first months of life, providing new insights into early neurodevelopment. Future studies should focus on establishing normative spatial patterns of functional network development across infancy and comparing them with neurodevelopmental disorders such as autism spectrum disorder (ASD). Identifying deviations in network structure and connectivity in high-risk infants could provide early biomarkers for ASD, facilitating early diagnosis and targeted interventions.

## Acknowledgment

This work was supported by NSF grant number 2112455 and NSF grant number 2316421.

## References

[1] A. N. Nielsen et al., “Maturation of large-scale brain systems over the first month of life,” Cerebral Cortex, vol. 33, pp. 2788–2803, 2023, doi: 10.1093/cercor/bhac242.

[2] V. J. Sydnor et al., “Neurodevelopment of the association cortices: Patterns, mechanisms, and implications for psychopathology,” Neuron, vol. 109, no. 18, pp. 2820–2846, Sep. 2021, doi: 10.1016/J.NEURON.2021.06.016.

[3] J. H. Gilmore, R. C. Knickmeyer, and W. Gao, “Imaging structural and functional brain development in early childhood,” Nat Rev Neurosci, vol. 19, no. 3, p. 123, Feb. 2018, doi: 10.1038/NRN.2018.1.

[4] A. N. Nielsen, D. J. Greene, C. Gratton, N. U. F. Dosenbach, S. E. Petersen, and B. L. Schlaggar, “Evaluating the Prediction of Brain Maturity From Functional Connectivity After Motion Artifact Denoising,” Cerebral Cortex, vol. 29, no. 6, pp. 2455–2469, Jun. 2019, doi: 10.1093/CERCOR/BHY117.

[5] S. L. Bressler and V. Menon, “Large-scale brain networks in cognition: emerging methods and principles,” Trends Cogn Sci, vol. 14, no. 6, pp. 277–290, Jun. 2010, doi: 10.1016/J.TICS.2010.04.004.

[6] M. Seraji, M. Mohebbi, A. Safari, and B. Krekelberg, “Multiple sclerosis reduces synchrony of the magnocellular pathway,” PLoS One, vol. 16, no. 8, p. e0255324, Aug. 2021, doi: 10.1371/JOURNAL.PONE.0255324.

[7] A. Abrol et al., “Developmental and aging resting functional magnetic resonance imaging brain state adaptations in adolescents and adults: A large N (>47K) study,” Hum Brain Mapp, vol. 44, no. 6, pp. 2158–2175, Apr. 2023, doi: 10.1002/HBM.26200.

[8] M. Seraji, C. A. Ellis, M. S. E. Sendi, R. L. Miller, and V. D. Calhoun, “Uncovering Effects of Schizophrenia upon a Maximally Significant, Minimally Complex Subset of Default Mode Network Connectivity Features,” 2024 46th Annual International Conference of the IEEE Engineering in Medicine and Biology Society (EMBC), pp. 1–4, Jul. 2024, doi: 10.1109/EMBC53108.2024.10782953.

[9] E. G. Duerden, D. Card, I. D. Lax, E. J. Donner, and M. J. Taylor, “Alterations in frontostriatal pathways in children born very preterm,” Dev Med Child Neurol, vol. 55, no. 10, pp. 952–958, Oct. 2013, doi: 10.1111/DMCN.12198.

[10] C. D. Smyser, A. Z. Snyder, and J. J. Neil, “Functional Connectivity MRI in Infants: Exploration of the Functional Organization of the Developing Brain,” Neuroimage, vol. 56, no. 3, p. 1437, Jun. 2011, doi: 10.1016/J.NEUROIMAGE.2011.02.073.

[11] W. Gao et al., “Functional Network Development During the First Year: Relative Sequence and Socioeconomic Correlations,” Cereb Cortex, vol. 25, no. 9, pp. 2919–2928, Sep. 2015, doi: 10.1093/CERCOR/BHU088.

[12] M. Eyre et al., “The Developing Human Connectome Project: typical and disrupted perinatal functional connectivity,” Brain, vol. 144, no. 7, pp. 2199–2213, Jul. 2021, doi: 10.1093/BRAIN/AWAB118.

[13] V. D. Calhoun, T. Adali, G. D. Pearlson, and J. J. Pekar, “A method for making group inferences from functional MRI data using independent component analysis,” Hum Brain Mapp, vol. 14, no. 3, pp. 140–151, 2001, doi: 10.1002/HBM.1048.

[14] L. Ma, S. Shultz, Z. Fu, M. Seraji, A. Iraji, and V. Calhoun, “Spontaneous Brain Dynamics Associated With Acceleration Of Long-term Functional Connectome In Postnatal Development,” bioRxiv, p. 2024.11.14.623615, Nov. 2024, doi: 10.1101/2024.11.14.623615.

[15] Q. Li et al., “Altered Functional Network Energy Across Multiscale Brain Networks in Preterm vs. Full-Term Subjects: Insights from the Adolescent Brain Cognitive Development (ABCD) Study,” bioRxiv, p. 2025.02.09.637316, Feb. 2025, doi: 10.1101/2025.02.09.637316.

[16] Q. Li, M. Seraji, V. D. Calhoun, and A. Iraji, “Complexity Measures of Psychotic Brain Activity In The FMRI Signal,” Proceedings of the IEEE Southwest Symposium on Image Analysis and Interpretation, pp. 9–12, 2024, doi: 10.1109/SSIAI59505.2024.10508702.

[17] Y. Du and Y. Fan, “Group information guided ICA for fMRI data analysis,” Neuroimage, vol. 69, pp. 157–197, Apr. 2013, doi: 10.1016/J.NEUROIMAGE.2012.11.008.

[18] Q. Li et al., “Deciphering Multiway Multiscale Brain Network Connectivity: Insights from Birth to 6 Months,” bioRxiv, p. 2025.01.24.634772, Jan. 2025, doi: 10.1101/2025.01.24.634772.

[19] M. Seraji, S. Shultz, Q. Li, Z. Fu, V. D. Calhoun, and A. Iraji, “Spatial development of brain networks during the first six postnatal months,” Communications Biology 2025 8:1, vol. 8, no. 1, pp. 1–18, Oct. 2025, doi: 10.1038/s42003-025-08913-z.

[20] S. N. Wood, “Generalized additive models: An introduction with R, second edition,” Generalized Additive Models: An Introduction with R, Second Edition, pp. 1–476, Jan. 2017, doi: 10.1201/9781315370279/GENERALIZED-ADDITIVE-MODELS-SIMON-WOOD.

[21] E. Damaraju, A. Caprihan, J. R. Lowe, E. A. Allen, V. D. Calhoun, and J. P. Phillips, “Functional connectivity in the developing brain: A longitudinal study from 4 to 9 months of age,” Neuroimage, vol. 84, pp. 169–180, Jan. 2014, doi: 10.1016/J.NEUROIMAGE.2013.08.038.

[22] A. S. Hodel, “Rapid Infant Prefrontal Cortex Development and Sensitivity to Early Environmental Experience,” Dev Rev, vol. 48, p. 113, Jun. 2018, doi: 10.1016/J.DR.2018.02.003.

[23] N. Kriegeskorte, R. Cusack, and P. Bandettini, “How does an fMRI voxel sample the neuronal activity pattern: compact kernel or complex spatiotemporal filter?,” Neuroimage, vol. 49, no. 3, p. 1965, Feb. 2010, doi: 10.1016/J.NEUROIMAGE.2009.09.059.

[24] E. Wenger, C. Brozzoli, U. Lindenberger, and M. Lövdén, “Expansion & Renormalization of Human Brain Structure During Skill Acquisition,” Trends Cogn Sci, vol. 21, no. 12, p. 930, Dec. 2017, doi: 10.1016/J.TICS.2017.09.008.

[25] A. Diamond, “Close interrelation of motor development and cognitive development and of the cerebellum and prefrontal cortex,” Child Dev, vol. 71, no. 1, pp. 44–56, 2000, doi: 10.1111/1467-8624.00117.

[26] C. J. Stoodley and J. D. Schmahmann, “Functional topography in the human cerebellum: a meta-analysis of neuroimaging studies,” Neuroimage, vol. 44, no. 2, pp. 489–501, Jan. 2009, doi: 10.1016/J.NEUROIMAGE.2008.08.039.

[27] U. A. Tooley, D. S. Bassett, and A. P. Mackey, “Environmental influences on the pace of brain development,” Nat Rev Neurosci, vol. 22, no. 6, p. 372, Jun. 2021, doi: 10.1038/S41583-021-00457-5.

[28] M. M. K. Bruchhage, G. C. Ngo, N. Schneider, V. D’Sa, and S. C. L. Deoni, “Functional connectivity correlates of infant and early childhood cognitive development,” Brain Struct Funct, vol. 225, no. 2, pp. 669–681, Mar. 2020, doi: 10.1007/S00429-020-02027-4/TABLES/3.

[29] W. Gao, W. Lin, K. Grewen, and J. H. Gilmore, “Functional Connectivity of the Infant Human Brain: Plastic and Modifiable,” The Neuroscientist, vol. 23, no. 2, p. 169, Apr. 2017, doi: 10.1177/1073858416635986.

[30] W. Lin et al., “Functional Connectivity MR Imaging Reveals Cortical Functional Connectivity in the Developing Brain,” AJNR Am J Neuroradiol, vol. 29, no. 10, p. 1883, Nov. 2008, doi: 10.3174/AJNR.A1256.

